## Supplemental Data for "WHIMP links the actin nucleation machinery to Src-family kinase signaling during protrusion and motility"

**Supplemental Figures and Legends**

| Description | Vector | Species | AA | R.E. Sites | Source |
|---|---|---|---|---|---|
| pKC-FastBac-MBP | pKC-FastBacMBP | N/A | N/A | N/A | Shen et al., 2012 |
| pKC-FastBac-MBP-WHAMM | pKC-FastBacMBP | Human | 1-809 | KpnI-NotI | Shen et al., 2012 |
| pKC-FastBac-MBP-WHIMP | pKC-FastBacMBP | Mouse | 1-516 | XhoI-HindIII | This study |
| pKC-FastBac-MBP-WASP(WCA) | pKC-FastBacMBP | Human | 405-502 | KpnI-NotI | This study |
| pKC-FastBac-MBP-N-WASP(WWCA) | pKC-FastBacMBP | Rat | 386-501 | KpnI-NotI | This study |
| pKC-FastBac-MBP-WAVE2(WCA) | pKC-FastBacMBP | Mouse | 402-497 | KpnI-NotI | This study |
| pKC-FastBac-MBP-WASH(WCA) | pKC-FastBacMBP | Human | 326-465 | KpnI-NotI | This study |
| pKC-FastBac-MBP-WHAMM(WWCA) | pKC-FastBacMBP | Human | 659-809 | KpnI-NotI | Shen et al., 2012 |
| pKC-FastBac-MBP-JMY(WWWCA) | pKC-FastBacMBP | Mouse | 817-983 | KpnI-NotI | This study |
| pKC-FastBac-MBP-WHIMP(WCA) | pKC-FastBacMBP | Mouse | 449-516 | KpnI-NotI | This study |
| pKC-EGFP-C1 (vector) | pKC-EGFP-C1 | N/A | N/A | N/A | Campellone et al., 2008 |
| pKC-EGFP-N-WASP(WWCA) | pKC-EGFP-C1 | Rat | 386-501 | KpnI-EcoRI | This study |
| pKC-EGFP-WHIMP(WCA) | pKC-EGFP-C1 | Mouse | 449-516 | KpnI-NotI | This study |
| pKC-EGFP-WHIMP | pKC-EGFP-C1 | Mouse | 1-516 | EcoRI-NotI | This study |
| pKC-mCherry-C1 (vector) | pKC-mCherry-C1 | N/A | N/A | N/A | Campellone et al., 2008 |
| pKC-mCherry-WHIMP | pKC-mCherry-C1 | Mouse | 1-516 | EcoRI-NotI | This study |
| pKC-LAP-C1 (vector) | pKC-LAP-C1 | N/A | N/A | N/A | Campellone et al., 2008 |
| pKC-LAP-WHIMP | pKC-LAP-C1 | Mouse | 1-516 | SpeI-EcoRI | This study |
| pKC-LAP-WHIMP(ΔWCA) | pKC-LAP-C1 | Mouse | 1-448 | SpeI-EcoRI | This study |
| pGFP (pKC425) (vector) | pCDNA3::GFP | N/A | N/A | N/A | Campellone et al., 2008 |
| pGFP-N-WASP | pCDNA3::GFP-Flag | Rat | 1-501 | KpnI-EcoRI | Campellone et al., 2008 |
| pGFP-Cortactin | pCDNA3::GFP | Mouse | 1-546 | EcoRI-BamHI | Campellone et al., 2008 |
| pmCherry-Rab5a | pmCherry-C1 | Mouse | | | Addgene #27679 |
| pmCherry-Rab5a(Q79L) | pmCherry-C1 | Human | | | Addgene #35138 |
| pEGFP-Rac1(Q61L) | pCDNA3-EGFP | Human | | | Addgene #12981 |

**Table S1**

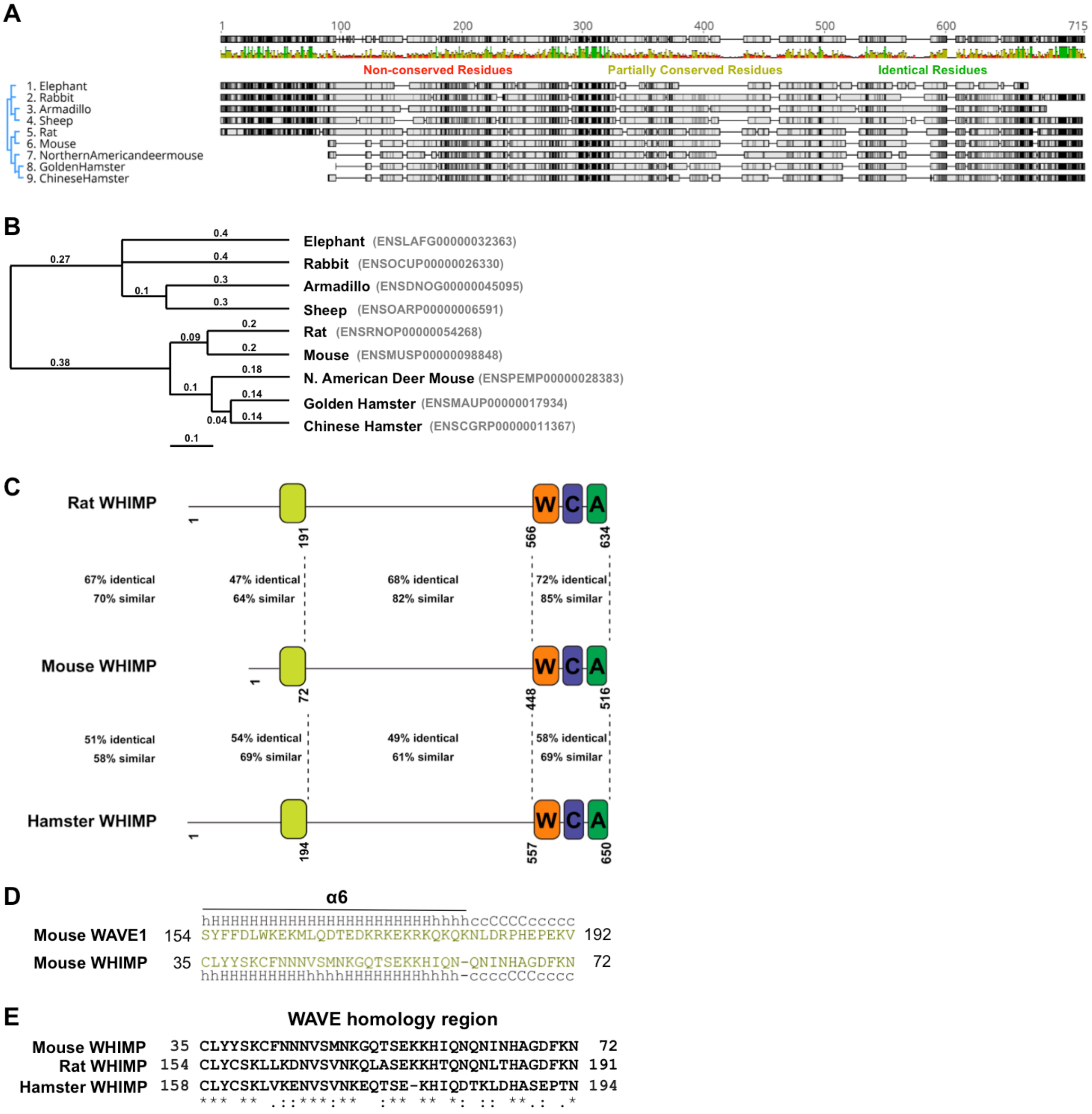
A
Non-conserved Residues
Partially Conserved Residues
Identical Residues
1. Elephant
2. Rabbit
3. Armadillo
4. Sheep
5. Rat
6. Mouse
7. NorthernAmericandeermouse
8. GoldenHamster
9. ChineseHamster
B
Elephant (ENSLAFG00000032363)
Rabbit (ENSOCUP00000026330)
Armadillo (ENSDNOG00000045095)
Sheep (ENSOARP00000006591)
Rat (ENSRNOP00000054268)
Mouse (ENSMUSP00000098848)
N. American Deer Mouse (ENSPEMP00000028383)
Golden Hamster (ENSMAUP00000017934)
Chinese Hamster (ENSCGRP00000011367)
C
Rat WHIMP
Mouse WHIMP
Hamster WHIMP
67% identical 70% similar
47% identical 64% similar
68% identical 82% similar
72% identical 85% similar
51% identical 58% similar
54% identical 69% similar
49% identical 61% similar
58% identical 69% similar
D
α6
hHHHHHHHHHHHHHHHHHHHHHHHHhhhhccCCCCccccc
Mouse WAVE1 154 SYFFDLWKEKMLQDTEDKRKEKRKQKQKNLDRPHEPEKV 192
Mouse WHIMP 35 CLYYSKCFNNNVSMNKGQTSEKKHIQN-QNINHAGDFKN 72
hhHHHHHHHHHhhhhHHHHHHHHHhhhh-ccccCCCcccc
E
WAVE homology region
Mouse WHIMP 35 CLYYSKCFNNNVSMNKGQTSEKKHIQNQNINHAGDFKN 72
Rat WHIMP 154 CLYCSKLLKDNVSVNKQLASEKKHTQNQNLTHAGDFKN 191
Hamster WHIMP 158 CLYCSKLVKENVSVNKEQTSE-KHIQDTKLDHASEPTN 194
*** ** .::***:** :** ** *: :: **.: .*

**Fig. S1**

**Fig. S1. WHIMP orthologs are present in multiple vertebrates. (A)** A consensus sequence illustration of WHIMP orthologs aligned using Geneious software is shown. Non-conserved regions are depicted in white, while partially-conserved and identical residues are shown in gray and black. **(B)** A phylogenetic tree of WHIMP was constructed using estimates of maximum likelihood phylogenies (PhyML) from alignments of amino acid sequences generated using Geneious software. UniProt ID numbers are listed next to the common animal names. The scale bar indicates the branch length and amino acid substitutions per site. **(C)** Primary domain organizations for mouse (*Mus musculus*), rat, and Chinese hamster WHIMP, along with their sequence identities and similarities from EMBOSS Needle are shown. **(D)** The HHpred algorithm predicts sequence similarity between amino acids 154-192 of the WAVE1 α6 helix and residues 35-72 of WHIMP. H/h=helix and C/c=coil, where uppercase letters represent higher confidence predictions. **(E)** Multiple sequence alignment of the α6 WAVE homology region of WHIMP from mouse, rat, and hamster is shown.

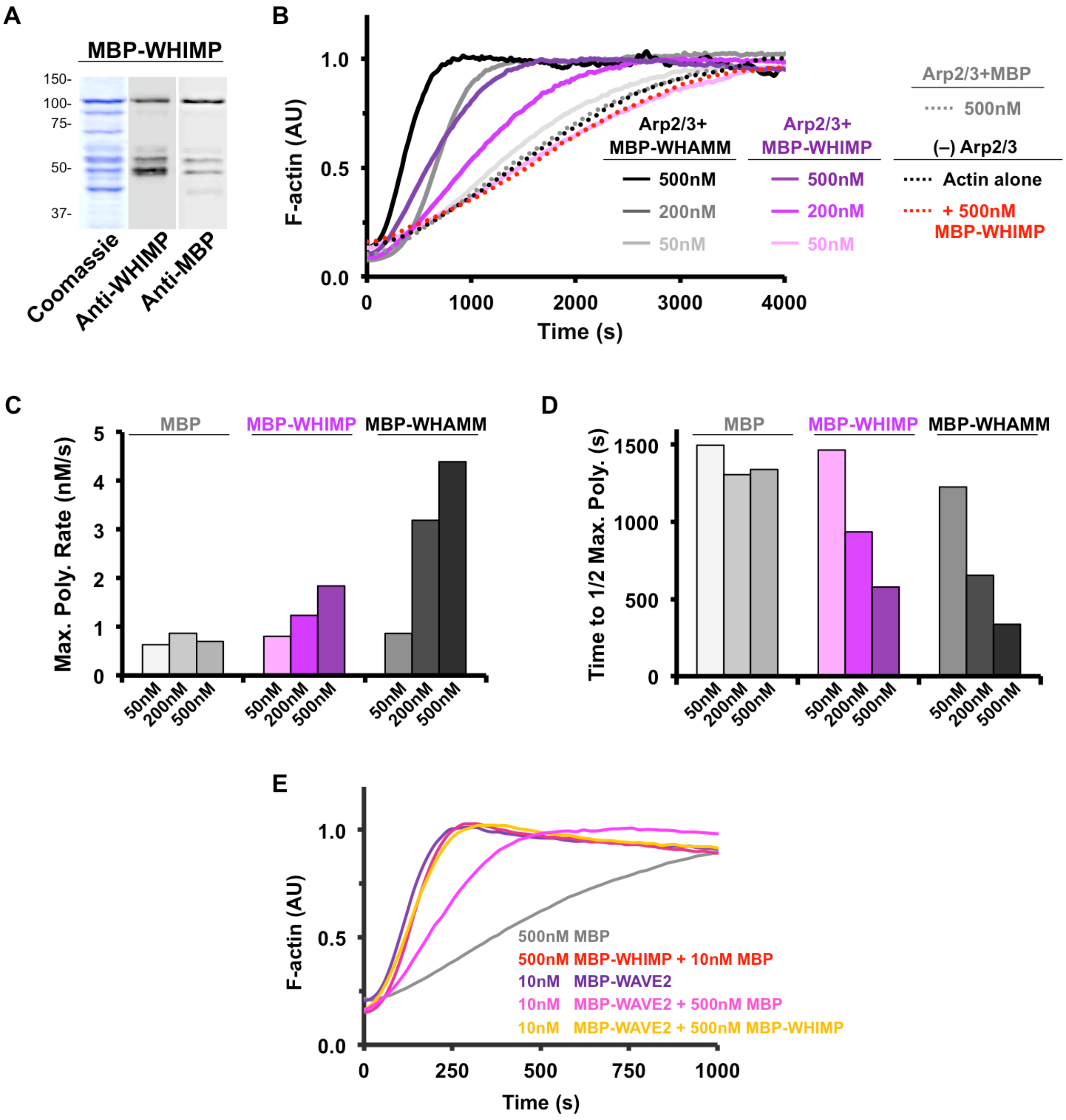
A
MBP-WHIMP
150-
100-
75-
50-
37-
Coomassie
Anti-WHIMP
Anti-MBP
B
F-actin (AU)
Time (s)
Arp2/3+ MBP-WHAMM
500nM
200nM
50nM
Arp2/3+ MBP-WHIMP
500nM
200nM
50nM
Arp2/3+MBP
500nM
(–) Arp2/3
Actin alone
+ 500nM MBP-WHIMP
C
Max. Poly. Rate (nM/s)
MBP
MBP-WHIMP
MBP-WHAMM
50nM
200nM
500nM
D
Time to 1/2 Max. Poly. (s)
MBP
MBP-WHIMP
MBP-WHAMM
E
F-actin (AU)
Time (s)
500nM MBP
500nM MBP-WHIMP + 10nM MBP
10nM MBP-WAVE2
10nM MBP-WAVE2 + 500nM MBP
10nM MBP-WAVE2 + 500nM MBP-WHIMP

**Fig. S2**

**Fig. S2. WHIMP is a weak actin nucleation-promoting factor *in vitro*. (A)** Purified maltose binding protein (MBP) tagged full-length WHIMP was subjected to SDS-PAGE followed by staining with Coomassie blue or immunoblotting with anti-WHIMP or anti-MBP antibodies. **(B)** Representative actin polymerization assays using 2μM actin, 20nM Arp2/3 complex, and the indicated concentrations of MBP, MBP-WHIMP, or MBP-WHAMM are shown. AU, arbitrary units. **(C)** Maximum actin polymerization rates were calculated from curves shown in panel B. **(D)** Times to half-maximal polymer were calculated from curves shown in panel B. **(E)** Actin polymerization assays using 2μM actin, 20nM Arp2/3 complex, and a combination of MBP, MBP-WHIMP(WCA), and MBP-WAVE2(WCA) are shown. AU, arbitrary units.

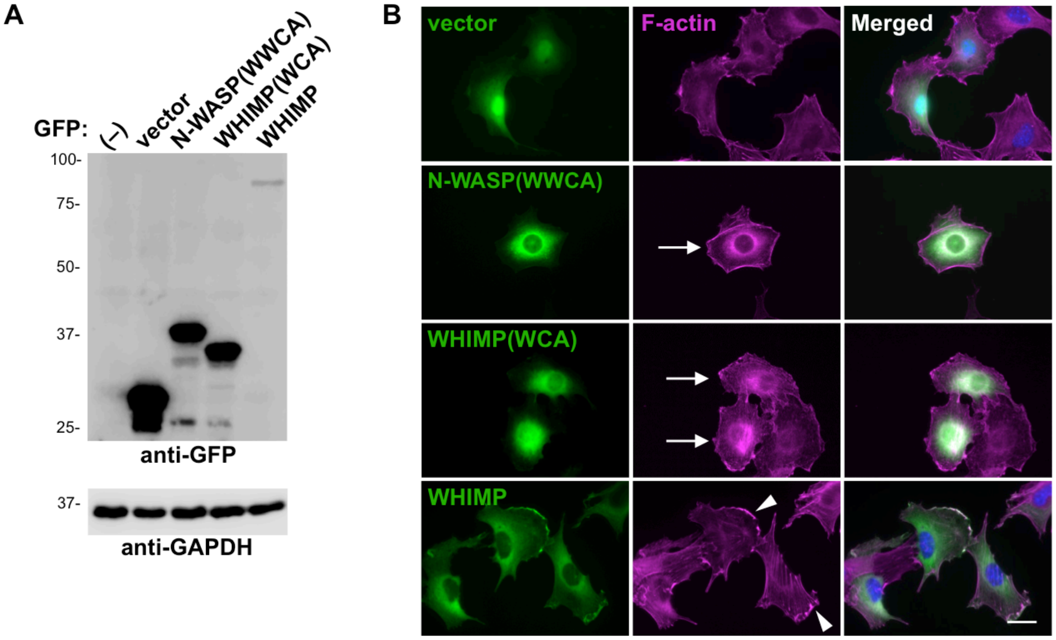
A
GFP:
(–)
vector
N-WASP(WWCA)
WHIMP(WCA)
WHIMP
100-
75-
50-
37-
25-
anti-GFP
37-
anti-GAPDH
B
vector
F-actin
Merged
N-WASP(WWCA)
WHIMP(WCA)
WHIMP

Fig. S3

**Fig. S3. WHIMP induces cytoplasmic actin assembly and localizes to membrane protrusions in B16-F1 cells. (A)** B16-F1 cells transiently transfected with plasmids encoding GFP or GFP-tagged N-WASP(WWCA), WHIMP(WCA), or full-length WHIMP were lysed and subjected to SDS-PAGE and immunoblotting with antibodies to GFP and GAPDH. **(B)** Cells treated as in A were fixed and stained with phalloidin to visualize F-actin (magenta) and DAPI to label DNA (blue). Arrows highlight GFP-WCA-expressing cells with increased cytoplasmic F-actin content, while arrowheads indicate prominent F-actin-rich protrusions in GFP-WHIMP-expressing cells. Scale bar, 20μm.

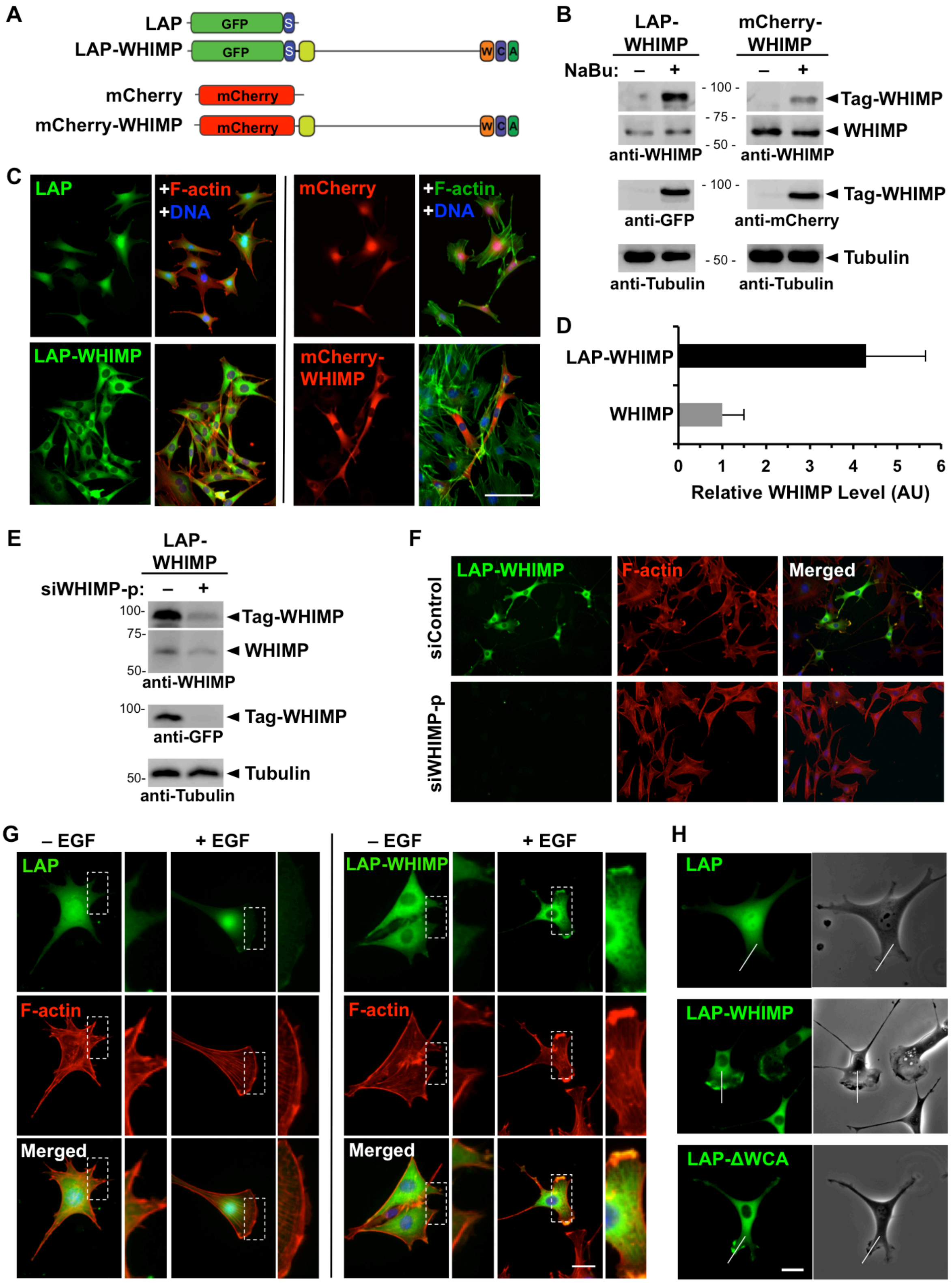
A
LAP
GFP
S
LAP-WHIMP
W
C
A
mCherry
mCherry-WHIMP
B
LAP-WHIMP
mCherry-WHIMP
NaBu:
– +
- 100 -
- 75 -
- 50 -
Tag-WHIMP
WHIMP
anti-WHIMP
anti-GFP
anti-mCherry
Tubulin
anti-Tubulin
C
LAP
+F-actin
+DNA
LAP-WHIMP
mCherry
mCherry-WHIMP
D
LAP-WHIMP
WHIMP
0 1 2 3 4 5 6
Relative WHIMP Level (AU)
E
LAP-WHIMP
siWHIMP-p:
100-
75-
50-
Tag-WHIMP
WHIMP
anti-WHIMP
anti-GFP
Tubulin
anti-Tubulin
F
siControl
siWHIMP-p
LAP-WHIMP
F-actin
Merged
G
– EGF
+ EGF
LAP
F-actin
Merged
LAP-WHIMP
H
LAP
LAP-WHIMP
LAP-ΔWCA

Fig. S4

**Fig. S4. NIH3T3 cell lines engineered to express fluorescently-tagged WHIMP show WHIMP localization to membrane protrusions. (A)** Diagrams of the fusion proteins used to generate fluorescent NIH3T3 cell lines are shown. The LAP-tag includes GFP and an S-peptide. **(B)** NIH3T3 cells stably encoding LAP-WHIMP or mCherry-WHIMP were treated with 10mM sodium butyrate (NaBu) for 16-18h and sorted into GFP- or mCherry-expressing populations using flow cytometry. Following outgrowth, cells were left untreated (–) or were treated (+) with NaBu, lysed, and subjected to SDS-PAGE and immunoblotting with antibodies to WHIMP, GFP, mCherry, or Tubulin. The tagged and endogenous versions of WHIMP are indicated. **(C)** Cells treated as in B were fixed and stained with phalloidin to visualize F-actin and DAPI to label DNA. Scale bar, 100μm. **(D)** Cells induced to express LAP-WHIMP were immunoblotted with antibodies to WHIMP. The mean level of LAP-WHIMP relative to endogenous WHIMP was determined following densitometry of 6 representative blots +/-SD. **(E)** Cells induced to express LAP-WHIMP were treated with either a control siRNA or a pool of WHIMP-specific siRNAs for 48h, lysed, and subjected to SDS-PAGE and immunoblotting for WHIMP, GFP, and Tubulin. **(F)** Cells treated as in E were fixed and stained with phalloidin to visualize F-actin and with DAPI to label DNA. **(G)** Cells stably encoding LAP or LAP-WHIMP were induced, serum-starved, and stimulated with EGF for 5min. Cells were then fixed and stained with phalloidin and DAPI. Magnifications highlight LAP-WHIMP-specific enrichment at the cell periphery and co-localization with F-actin. Scale bar, 20μm. **(H)** Cells stably encoding LAP, LAP-WHIMP, or LAP-WHIMP(ΔWCA) were induced and examined live for GFP fluorescence and by phase-contrast microscopy. White lines indicate positions in which kymograph analyses were performed in Fig.5I. Scale bar, 20μm.

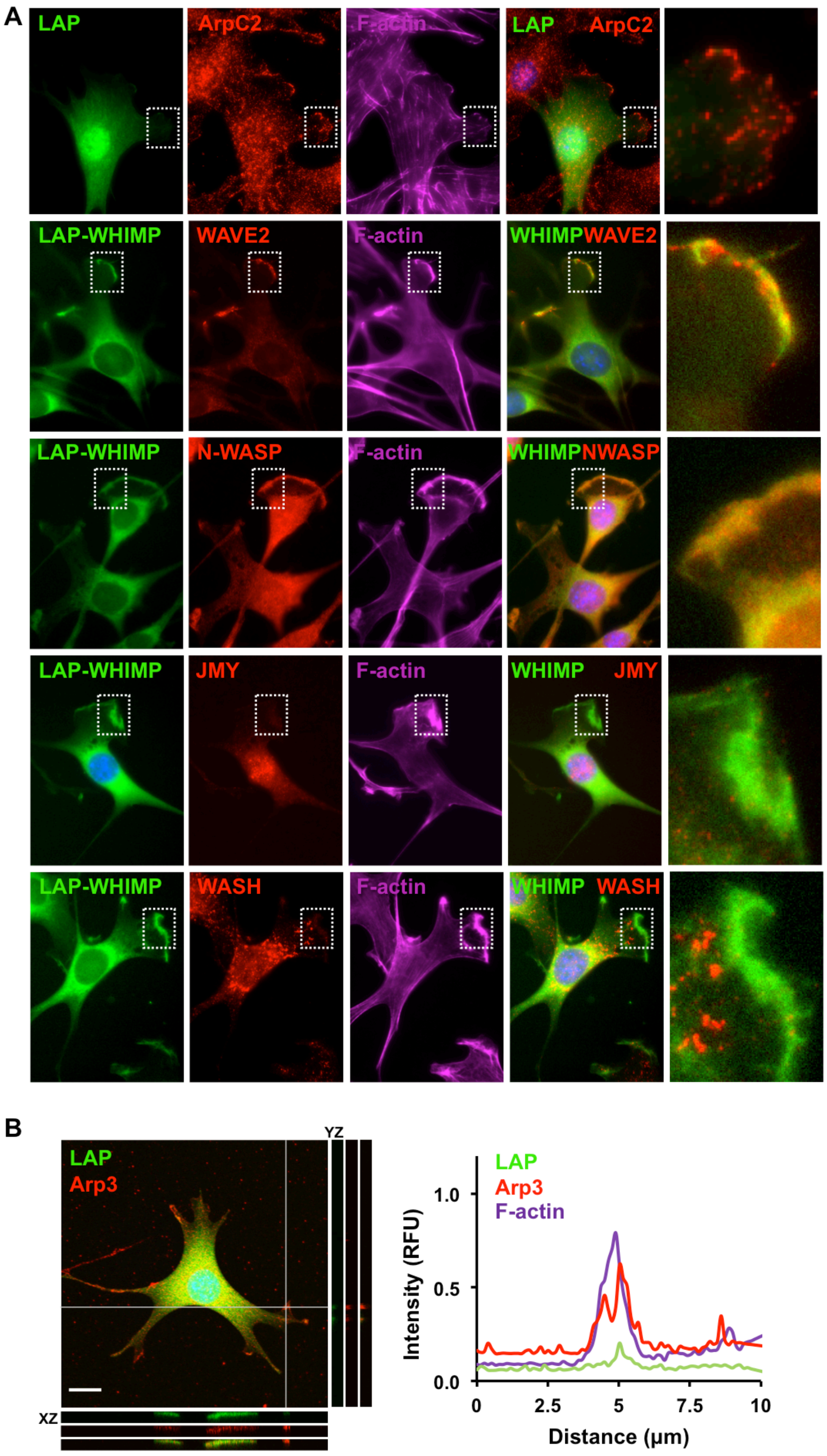
A
LAP
ArpC2
F-actin
LAP ArpC2
LAP-WHIMP
WAVE2
F-actin
WHIMP WAVE2
LAP-WHIMP
N-WASP
F-actin
WHIMP NWASP
LAP-WHIMP
JMY
F-actin
WHIMP JMY
LAP-WHIMP
WASH
F-actin
WHIMP WASH
B
LAP
Arp3
YZ
XZ
LAP
Arp3
F-actin
Intensity (RFU)
1.0
0.5
0.0
0
2.5
5
7.5
10
Distance (µm)

**Fig. S5**

**<u>Fig. S5</u>. WAVE2 and N-WASP, but not JMY or WASH, localize to WHIMP-associated membrane protrusions. (A)** NIH3T3 cell lines expressing a LAP tag or LAP-WHIMP were fixed and stained with antibodies to ArpC2, WAVE2, N-WASP, JMY, or WASH (red), phalloidin to visualize F-actin (magenta), and DAPI to label DNA (blue). Magnifications (right column) highlight LAP-WHIMP-specific localization to ruffles and overlap with WAVE2 and N-WASP. **(B)** LAP-expressing cells stained as in A were subjected to confocal microscopy for comparison to Fig.4D. Maximum intensity projections are shown in the large panel, and adjacent orthogonal views of YZ and XZ planes are indicated by the two gray lines. The plot profiles depict pixel intensities of LAP, Arp3, and F-actin along a 10μm line drawn near the intersection of the two gray lines. Scale bars, 20μm.

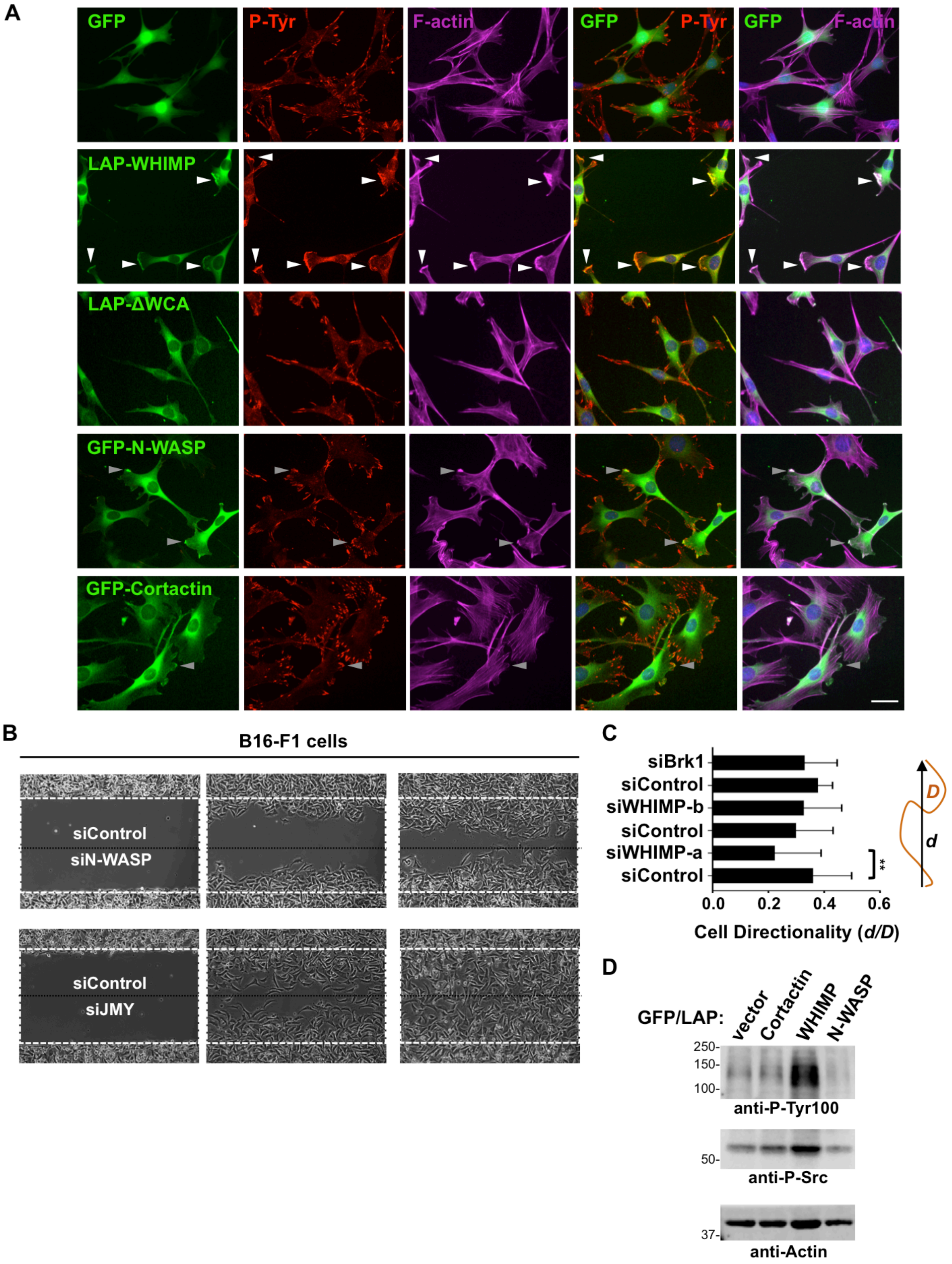
A
GFP
P-Tyr
F-actin
GFP P-Tyr
GFP F-actin
LAP-WHIMP
LAP-ΔWCA
GFP-N-WASP
GFP-Cortactin
B
B16-F1 cells
siControl
siN-WASP
siControl
siJMY
C
siBrk1
siControl
siWHIMP-b
siControl
siWHIMP-a
siControl
**
0.0
0.2
0.4
0.6
Cell Directionality (d/D)
D
d
D
GFP/LAP:
vector
Cortactin
WHIMP
N-WASP
250-
150-
100-
anti-P-Tyr100
50-
anti-P-Src
37-
anti-Actin

**Fig. S6**

**<u>Fig. S6</u>. WHIMP-associated membrane ruffling, tyrosine phosphorylation, and cell motility phenotypes are distinct from those associated with N-WASP or Cortactin. (A)** NIH3T3 cells stably encoding GFP, LAP-WHIMP, LAP-WHIMP(ΔWCA), GFP-N-WASP, or GFP-Cortactin were treated with sodium butyrate for 16-18h, fixed, and stained with antibodies to detect phosphotyrosines (red), phalloidin to visualize F-actin (magenta), and DAPI to label DNA (blue). White arrowheads highlight large WHIMP- and F-actin-associated membrane ruffles, while grey arrowheads point to occasional small protrusions in N-WASP- or Cortactin-overexpressing cells. Scale bar, 20μm. **(B)** B16-F1 cells were treated with siRNAs, grown in cell reservoirs separated by a 0.5mm barrier, and analyzed by phase-contrast microscopy at 5min intervals for 12h after removing the barrier. Control cells are shown in the top chamber, and nucleation factor-depleted cells are shown in the bottom. The white dashed box encompasses the wound area at t=0, and the black dashed line indicates the midpoint of the cell-free area. **(C)** A directionality index was calculated as the ratio of the distance between the starting and ending point "*d*" and the trajectory "*D*" based on the cell tracks described in Fig.7E. The bars represent the mean directionality +/-SD of cells (n≥20) for each siRNA treatment. **p<0.01 (two-tailed t-test). **(D)** Cells induced to express GFP, GFP-Cortactin, LAP-WHIMP, or GFP-N-WASP were subjected to SDS-PAGE and immunoblotting with antibodies to phosphotyrosine, active Src P-Tyr416 (P-Src), and actin.

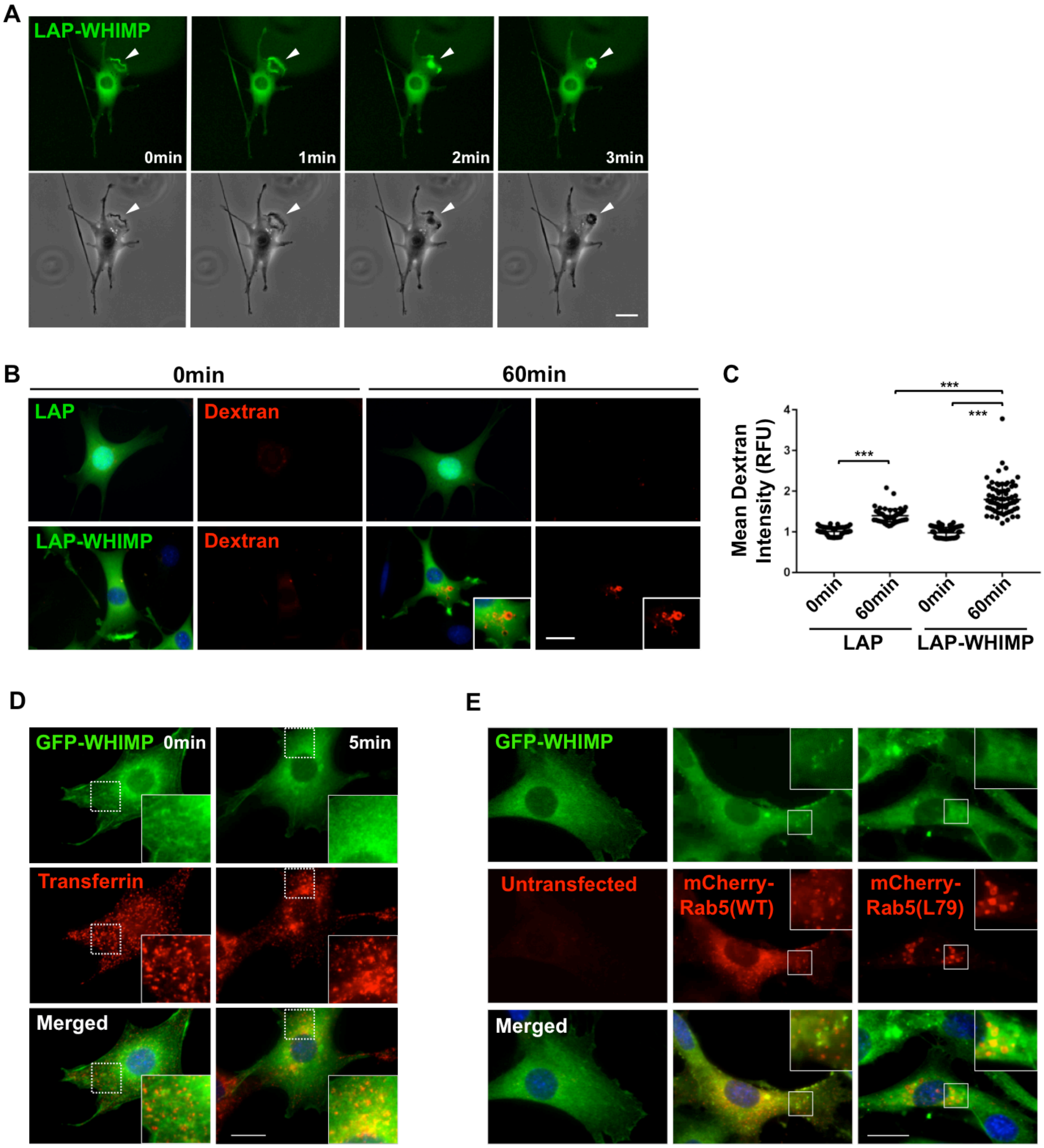
A
LAP-WHIMP
0min
1min
2min
3min
B
0min
60min
LAP
Dextran
LAP-WHIMP
Dextran
C
Mean Dextran Intensity (RFU)
***
***
***
0min
60min
0min
60min
LAP
LAP-WHIMP
D
GFP-WHIMP
0min
5min
Transferrin
Merged
E
GFP-WHIMP
Untransfected
mCherry-Rab5(WT)
mCherry-Rab5(L79)
Merged

Fig. S7

<u>Fig. S7</u>. **Macropinocytosis, but not receptor-mediated endocytosis, is enhanced by WHIMP overexpression.** **(A)** GFP fluorescence and phase-contrast timelapse images of live LAP-WHIMP-expressing NIH3T3 cells are shown. Arrowheads highlight dorsal ruffle formation and circular constriction. **(B)** Cells expressing LAP or LAP-WHIMP were incubated with 70kDa TMR-dextran (red) for 0 or 60min, fixed, and stained with DAPI to label DNA (blue). Insets highlight increased amounts of TMR-dextran in LAP-WHIMP cells after the 60min incubation. **(C)** The mean TMR-dextran intensity per cell was measured using ImageJ. Each point represents the mean intensity of a single cell ($n \geq 30$) compiled from multiple experiments, and the horizontal line represents the overall mean +/- SD. **(D)** NIH3T3 cells were transiently transfected with a plasmid encoding GFP-WHIMP, and after 24h incubated with Alexa568-transferrin (10µg/ml) at 4°C in serum starvation media. Cells were then shifted to 37°C for 5min, fixed, and stained with DAPI. Insets highlight a lack of co-localization of GFP-WHIMP with punctate transferrin. **(E)** Cells were transiently co-transfected with plasmids encoding GFP-WHIMP and either wild type Rab5a or constitutively active Rab5a(Q79L), and after 24h fixed and stained with DAPI. Insets highlight a lack of co-localization between diffuse GFP-WHIMP and either wild type Rab5 or large endosome-associated Rab5(Q79L). *** $p<0.001$ (ANOVA with Tukey post-test). Scale bars, 20µm.

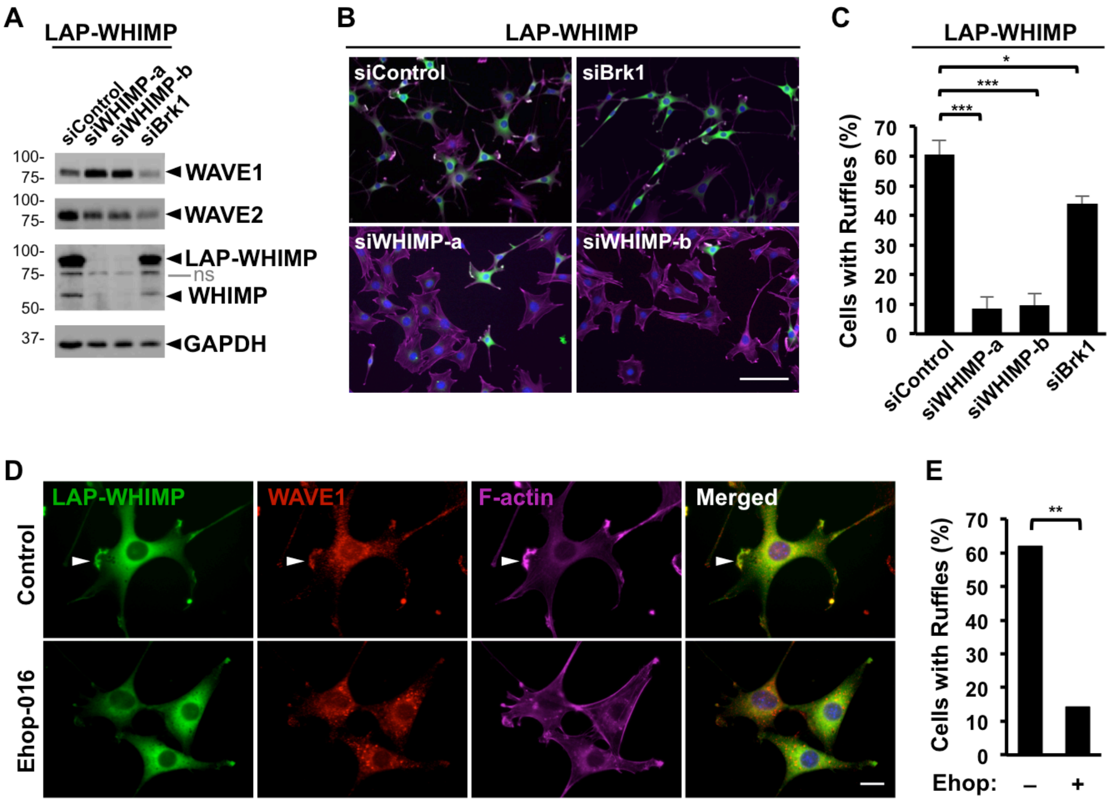
A
LAP-WHIMP
siControl
siWHIMP-a
siWHIMP-b
siBrk1
100-
75-
WAVE1
WAVE2
LAP-WHIMP
ns
WHIMP
50-
37-
GAPDH
B
LAP-WHIMP
siControl
siBrk1
siWHIMP-a
siWHIMP-b
C
LAP-WHIMP
Cells with Ruffles (%)
*
***
***
0
10
20
30
40
50
60
70
siControl
siWHIMP-a
siWHIMP-b
siBrk1
D
LAP-WHIMP
WAVE1
F-actin
Merged
Control
Ehop-016
E
**
Cells with Ruffles (%)
Ehop:
–
+

Fig. S8

**<u>Fig. S8</u>. WHIMP-induced membrane ruffling is partially dependent on Rac/WAVE activities. (A)** NIH3T3 cells stably-encoding LAP-WHIMP were treated with a control siRNA, independent siRNAs to WHIMP, or an siRNA to Brk1, induced to express the fusion protein, lysed, subjected to SDS-PAGE, and immunoblotted with antibodies to WAVE1, WAVE2, WHIMP, and GAPDH. The tagged and endogenous versions of WHIMP are indicated. ns, non-specific. **(B)** LAP-WHIMP cells were treated with siRNAs, fixed, and stained with phalloidin (magenta), and DAPI (blue). Scale bar, 100μm. **(C)** The % of cells with ruffles was quantified. Each bar represents the mean % of GFP-positive cells (n=200-300 per condition) with ruffles from 2-3 experiments +/-SD. *** $p < 0.001$, * $p < 0.05$ (ANOVA with Tukey post-test). **(D)** LAP-WHIMP-expressing cells were treated with 4μM EHop-016 for 5h, fixed, and stained with an antibody to WAVE1 (red), phalloidin (magenta), and DAPI (blue). The arrowhead highlights the position of a prominent WHIMP-associated ruffle. Scale bar, 20μm. **(E)** Each bar represents the % of cells (n=105-150 per condition) with ruffles from experiments performed in part D. ** $p<0.01$ (Fisher's exact test).

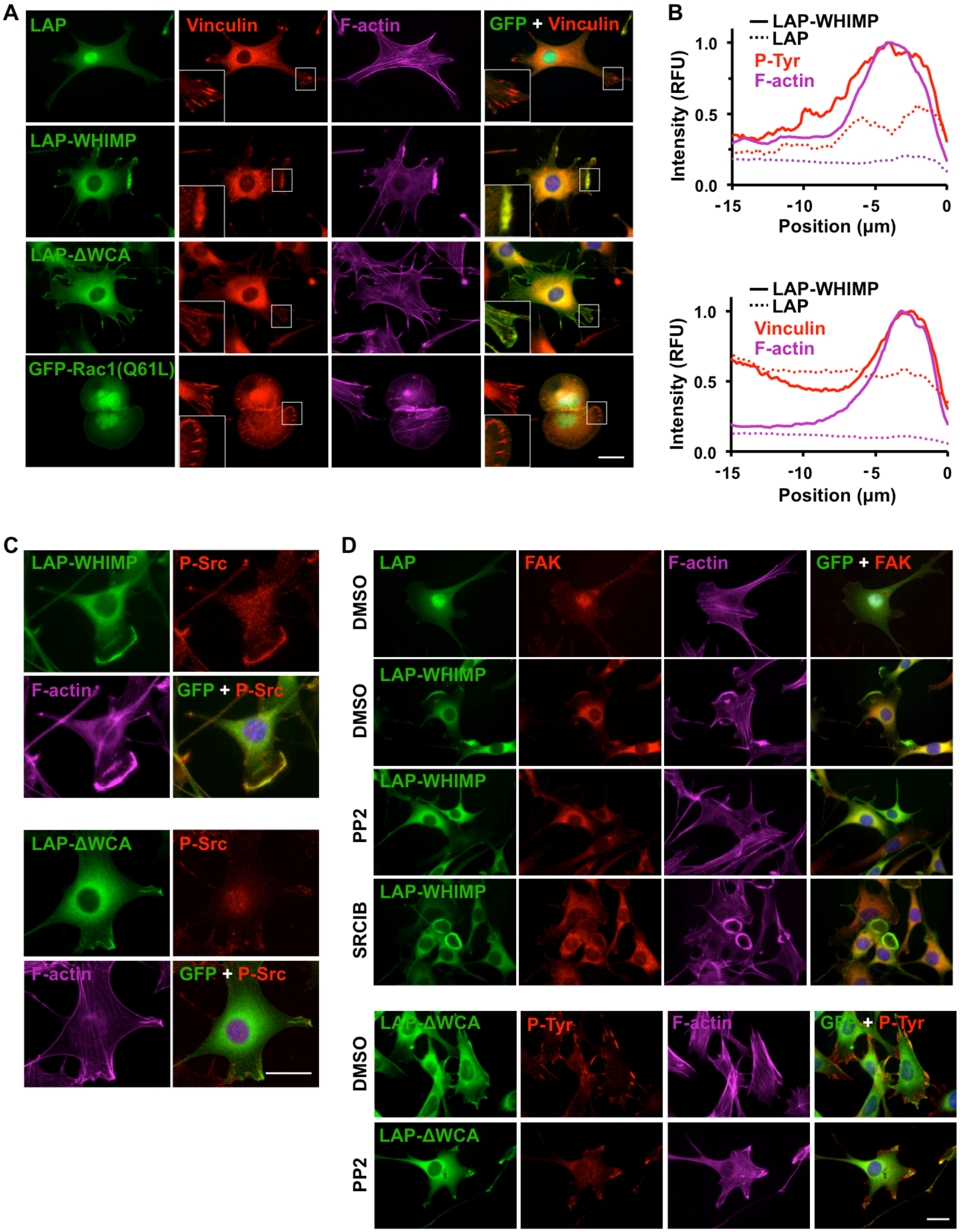
A
LAP
Vinculin
F-actin
GFP + Vinculin
LAP-WHIMP
LAP-ΔWCA
GFP-Rac1(Q61L)
B
LAP-WHIMP
LAP
P-Tyr
F-actin
Intensity (RFU)
Position (µm)
LAP-WHIMP
LAP
Vinculin
F-actin
Intensity (RFU)
Position (µm)
C
LAP-WHIMP
P-Src
F-actin
GFP + P-Src
LAP-ΔWCA
P-Src
F-actin
GFP + P-Src
D
DMSO
LAP
FAK
F-actin
GFP + FAK
DMSO
LAP-WHIMP
PP2
LAP-WHIMP
SRCIB
LAP-WHIMP
DMSO
LAP-ΔWCA
P-Tyr
F-actin
GFP + P-Tyr
PP2
LAP-ΔWCA

Fig. S9

**Fig. S9. Focal adhesion proteins are recruited to WHIMP-induced Src-dependent ruffles.** **(A)** NIH3T3 cell lines expressing LAP, LAP-WHIMP, or LAP-WHIMP(ΔWCA), or transiently transfected with a plasmid encoding GFP-Rac1(Q61L) were fixed and stained with antibodies to vinculin (red), phalloidin (magenta), and DAPI (blue). Insets highlight LAP-WHIMP localization at ruffles and broad enrichment of vinculin staining compared to wedge-like vinculin staining in other cells. **(B)** Line-scan plots depict the mean pixel intensity of phosphotyrosine (e.g., from Fig. 8A), vinculin, and F-actin staining along 15μm lines near the edges of LAP and LAP-WHIMP cells (n=10). The edges of the cells were set to 0μm on the X-axes. **(C)** Cell lines expressing LAP-WHIMP or LAP-WHIMP(ΔWCA) were fixed and stained with antibodies to active Src P-Tyr416 (P-Src; red), phalloidin (magenta), and DAPI (blue). **(D)** Cell lines expressing LAP, LAP-WHIMP, or LAP-WHIMP(ΔWCA) were treated with DMSO or with 20μM PP2 or Saracatinib (SRCIB) for 2h, fixed, and stained with antibodies to FAK or phosphotyrosine (red), phalloidin (magenta), and DAPI (blue). Scale bars, 20μm.

**Video S1. Timelapse phase-contrast movie of siControl (top) vs siWHIMP-a (bottom) wound closure (see Fig.7).**

**Video S2. Timelapse phase-contrast movie of siControl (top) vs siWHIMP-b (bottom) wound closure (see Fig.7).**

**Video S3. Timelapse phase-contrast movie of siControl (top) vs siBrk1 (bottom) wound closure (see Fig.7).**
